# Mechanical regulation of tissue flatness in *Marchantia*

**DOI:** 10.1101/2024.10.25.620180

**Authors:** Jordan Ferria, Carla J.A. Fournié, Magdalena H. Jankowska, Doron Grossman, Adrienne H.K. Roeder, Stéphanie Drevensek, Arezki Boudaoud

## Abstract

The production of flat organs involves a tight regulation of tissue expansion. During growth, the cell wall is under mechanical tension and continuous remodelling, requiring mechanisms to sense and maintain cell wall integrity (CWI). Here we studied the link between CWI sensing and flatness of *Marchantia polymorpha* thalli. We considered *FERONIA* (*FER*), a CWI sensor, and assessed its role in the maintenance of thallus shape. We developed a pipeline to quantify flatness at multiple scales. We found that both abolishing and increasing *FER* expression leads to loss of flatness. Based on experiments that reduce cell wall tension and partially restore wild-type cell size and thallus morphology, we ascribe loss of flatness to under- or to over-reaction of plants to mechanical stress. Altogether, our results support the concept that responses to mechanical stress contribute to the robustness of morphogenesis

## Introduction

Flat organs have evolved multiple times in numerous living organisms from flatworms to insect wings and plants. In plants, flat leaves have possibly evolved as a way to enhance light exposure and gas exchanges. The development of such a shape appears to require spatial regulation of cell proliferation and expansion because mutants disrupting these processes also commonly disrupt leaf flatness. For example, increased leaf curvature was observed in leaves of the *A. thaliana* mutants of the *PEAPOD2* (*PPD2)* and *NOVEL INTERACTOR OF JAZZ* (*NINJA*) genes, via the modulation of the position of the primary arrest front that separates cell proliferation dominated and and cell expansion dominated regions in leaves (Baekelandt et al., 2018; Du et al., 2018). Analysis of the *jaw-D* mutant also shows abnormal buckling phenotype near the leaf margin as a result of abnormal cell growth and division patterns (Palatnik et al., 2003; Harline et al., 2022). Similarly, loss of flatness was observed in the *angustifolia-1* (*an-1*) mutant of *Marchantia polymorpha* (Furuya et al., 2018). The *an-1* mutant displays twisting of the thallus (photosynthetic vegetative organ of *Marchantia polymorpha* corresponding to the gametophyte) along the growth axis, and a curled and wavy margin at the antheridiophores (mushroom-shaped organ containing male gametes). This alteration of flatness was suggested to be the result of abnormal orientation of cell expansion (Furuya et al., 2018), possibly through the regulation of cortical microtubule orientation as described in the *angustifolia* mutants of *Arabidopsis thaliana* (Folkers et al., 2002). The Marchantia mutants of *auxin response factor1* (*arf1*) also show alteration of thallus flatness and alteration of cell growth and division patterns, via alteration of the patterning of developmental axes by auxin (Kato et al., 2017; Flores-Sandoval et al., 2015). In *Arabidopsis thaliana,* ectopic expression mutants of the *ASYMMETRIC LEAVES 2* gene resulted in buckling of the abaxial sepals (Yadav et al., 2024), leading to variations in sepal thickness unlike in other studies where organ thickness was roughly constant in mutant plants. Together, this shows that organ flatness may rely on transcription factors and hormone gradients to regulate cell proliferation and expansion patterns.

Growth results from cell proliferation and expansion. Analysis of spatial variations of growth in petunia and tobacco leaves shows that flat leaf shape requires coordinated growth patterns (Mitchison et al., 2016, Alim et al., 2016). Indeed, growth gradients along a thin organ such as a leaf or a petal may result in conflicting tissue expansion (Coen et al., 2016) and induce mechanical stress in the tissue as cells with higher growth rates push on their neighbours (Boudaoud 2010). Such conflicts might be resolved by buckling: the loss of flatness and acquisition of a wavy shape due to compressive mechanical stress. Therefore, the development of flat shape requires tight regulation of growth to avoid buckling or folding (Boudaoud 2010, Palatnik et al., 2003; Sharon et al., 2004, Nath et al., 2003, Yadav et al., 2023, Harline et al., 2022). More specifically, heterogeneity in growth must be maintained below a certain threshold to avoid buckling and the production of fractal-like 3D shapes (Sharon et al., 2002; Audoly & Boudaoud 2003; Dervaux and Ben Amar, 2008; Efrati et al. 2009; Gemmer and Venkataramani 2013), like those of the leaves of ornamental cabbage.

The maintenance of flatness in growing organs suggests that growth heterogeneity is regulated. It was proposed that responses to mechanical stress induced by such heterogeneity may control growth and make it more homogeneous (Shraiman et al., 2005). In *Arabidopsis* sepals, cortical microtubules reorient circumferentially in cells around fast growing trichomes, along predicted stress patterns (Hervieux et al., 2017), possibly restraining trichome growth by guiding cellulose deposition and reinforcing cell walls circumferentially. In the *Arabidopsis* shoot apical meristem, *KATANIN* mediates alignment of cortical microtubules in response to mechanical stress and modulates the level of local cell-to-cell growth heterogeneity (Uyttewaal et al., 2012). Interestingly, Arabidopsis mutants of the *DEFECTIVE KERNEL1* (*DEK1*) gene are defective in activation of a mechanosensitive Ca2+ ion channel (Tran et al., 2017) while showing crinkled leaves (Johnson et al., 2008). Similarly, reduction of the touch response in the *vernalisation independence 3* (*vip3)* mutants were also associated with crinkled leaves (Trinh et al., 2024; Jensen et al., 2016).. Altogether, the perception of mechanical stress appears as a good candidate mechanism to coordinate cell growth, avoid buckling and maintain flatness.

The transmembrane protein FERONIA (FER) of the *Catharanthus roseus* RLK1-like (CrRLK1-like) receptor kinase subfamily was shown to prevent brassinosteroid induced cell growth, possibly as a way to prevent cell burst, consistent with its role in the maintenance of cell wall integrity (Höfte et al., 2015). *feronia* mutants also showed alteration of cell size and cell wall integrity upon salt stress, confirming a possible role in the maintenance of cell wall integrity (Feng et al., 2018). *feronia* mutants of *Arabidopsis thaliana* were found to have alteration in the calcium ion flux upon mechanical stimuli (Shih et al., 2014), indicating a role of *FERONIA* in responses to mechanical stress. The FER protein was also shown to play a significant role in modulating the penetration capacity of roots into the soil by sensing the relative hardness of the medium through activation of the transcription factor PIF3 and the mechanosensing ion channel PIEZO (Xu et al., 2024). Additionally, the capacity to promote microtubule rearrangement upon mechanical stress generated by single cell laser ablation, appears altered in the *fer* mutants (Tang et al., 2022), despite the presence of FERONIA independent mechanisms promoting cortical microtubule rearrangements (Malivert et al., 2024). Together, this suggests that FERONIA is involved in mechanoperception and mechanical stress response and that it plays a role in the sensing of mechanical stress induced by neighbouring cells. Given that FERONIA is a transmembrane protein, and that it can bind pectin via extracellular Malectin-A (MALA) domains (Feng et al., 2018), it can be hypothesised that FERONIA can detect variations in the tension of the cell wall and thus behave as a mechanosensor. Complementary experiments also showed that FER preferentially binds demethylesterified pectins (Lin et al., 2018), demonstrating that the FER protein can detect the chemical state of pectin. Furthermore, it was also shown that pectin binding is important for the acquisition of pavement cell shape in *A. thaliana* (Lin et al., 2022), demonstrating the necessity of a physical link to cell wall for complete protein activity. It was also suggested that the sensing of different states of pectin might be a common mechanism for the regulation of polar cell expansion (Lin et al., 2018), which indicates the crucial role of FER in touch perception. The Rapid Alkalinization Factor (RALF) ligand, a 49 amino acid peptide (Ryan and Pearce, 2004) was found to bind pectin and to promote phase separation of pectin fragments to induce Pectin-RALF-FERONIA-LORELEI clustering and trigger stress response (Liu et al., 2023). This suggests that FERONIA is involved in the detection of pectin fragments, suggesting a possible mechanism by which FERONIA could be involved in the surveillance of cell wall integrity. Over-expression of *AtRALF-ligand* also resulted in smaller root cells. Together, this suggests that RALF mediated activation of FERONIA could prevent cell expansion, possibly to prevent further mechanical stress in the context of compromised cell wall integrity (Liao et al., 2017). In *Marchantia polymorpha*, a single homologous gene of Arabidopsis *AtFER* was identified (Honkanen et al., 2016). Cellular force microscopy measurement of stiffness on the *fer-2* gemmae (small clonal structure involved in asexual reproduction) also suggests either a softer cell wall or lower turgor pressure. Softer cell wall would explain the presence of burst rhizoid cells observed in *fer-2* and *fer-3* mutants and the observation of dead cells in trypan blue exclusion tests (Mecchia et al., 2022). This suggests that *MpFERONIA* plays a similar role to that of *AtFERONIA* in the surveillance and maintenance of cell wall integrity. The *fer* mutants of *Marchantia* showed visible alteration of flatness, smaller cells and lower Young’s moduli (Mecchia et al., 2022). Here we consider flatness in relation with *FER* gene expression level and mechanical stress.

Three-dimensional reconstructions of *Marchantia polymorpha* thallus surface using micro-computed tomography have provided qualitative observation of the overall surface shape of flatness mutants such as *angustifolia-1* (Furuya et al., 2018). Despite satisfactory qualitative observations of flatness, the production of quantitative metrics has proven difficult and often relies on the measurement of curvature (Armon et al., 2014). While the Gaussian curvature efficiently describes curvature in a single point, it does not efficiently account for the variations of curvature at different scales. Multiple methods were developed in an attempt to better describe the complex curvature of leaves and their variations. The directional curvature of a leaf or curvature index can be approximated by computing the ratio of the absolute distance of the leaf tip from the base and the distance following the leaf (Liu et al., 2010; Ren et al., 2018). While this metric may broadly describe the overall leaf curvature, it lacks resolution along the leaf. The buckling of the leaf can be described as the perimeter to surface ratio (Nath et al., 2003) or as the proximodistal and mediolateral curvatures indexes taken together (Natarajan et al., 2020), but these methods do not account for the type of buckling that can be observed. While all the above methods provide sufficient metrics for a specific type of curvature or flatness defect, they fail to account for surface curvature as they cannot be used in the absence of a preferential axis of curvature such as observed in *Marchantia polymorpha.* Another method was developed that relies on the computation of the convex hull and overall surface as well as the convex hull and overall volume of the thalli (Furuya et al., 2019). This method enables efficient assessment of the volume spread of a thallus within space and proved efficient in discriminating the thallus shape of *angustifolia* mutants of *M. polymorpha* (Furuya et al., 2019). However, the convex score may be difficult to interpret in the context of flatness analysis, thus creating the need for new specific metrics. In this paper, we developed new methods to quantify flatness at multiple scales and investigated the alteration of flatness in the *fer-2* mutant and the *FERONIA* over-expressor (*FER-OE*) transgenic line of *Marchantia polymorpha* in an effort to identify the relevance of mechanosensing in the development of flat organs. We specifically looked at their response under osmotic treatments and the application of external mechanical constraints.

## Results

### Alteration of *FERONIA* expression induces a multi-scale loss of thallus flatness

We first asked whether the *FERONIA* (*FER*) gene has an influence on thallus flatness. We noticed that the *fer-2* mutant and the *FERONIA* over-expressor (*FER-OE*; _pro_*MpEF1::MpFERONIA-mCitrine*) transgenic line show an alteration of the overall thallus shape (fig. **1a**). We therefore used 3-dimensional micro computed tomography (μCT) to image plants, revealing a defect in the maintenance of the thallus flatness in those lines. The *fer-2* mutant exhibits an overall alteration of flatness and a tortuous shape while the over expression transgenic line shows alteration of flatness and hyponastic behaviour (fig. **1b**). From these µCT scans a flatness score was computed. The ϕ flatness score consists of the standard deviation of the thallus surface along the z-axis divided by the square root of the thallus projected area. High values indicate a lack of flatness, and a flat surface has a ϕ score of ‘0’. The *fer-2* mutant and *FER-OE* transgenic line show mean values of ϕ three times higher (0.18) than that of the *wild type* (0.06) confirming a consistent defect in flatness (fig. **1c**).

**Figure 1:**
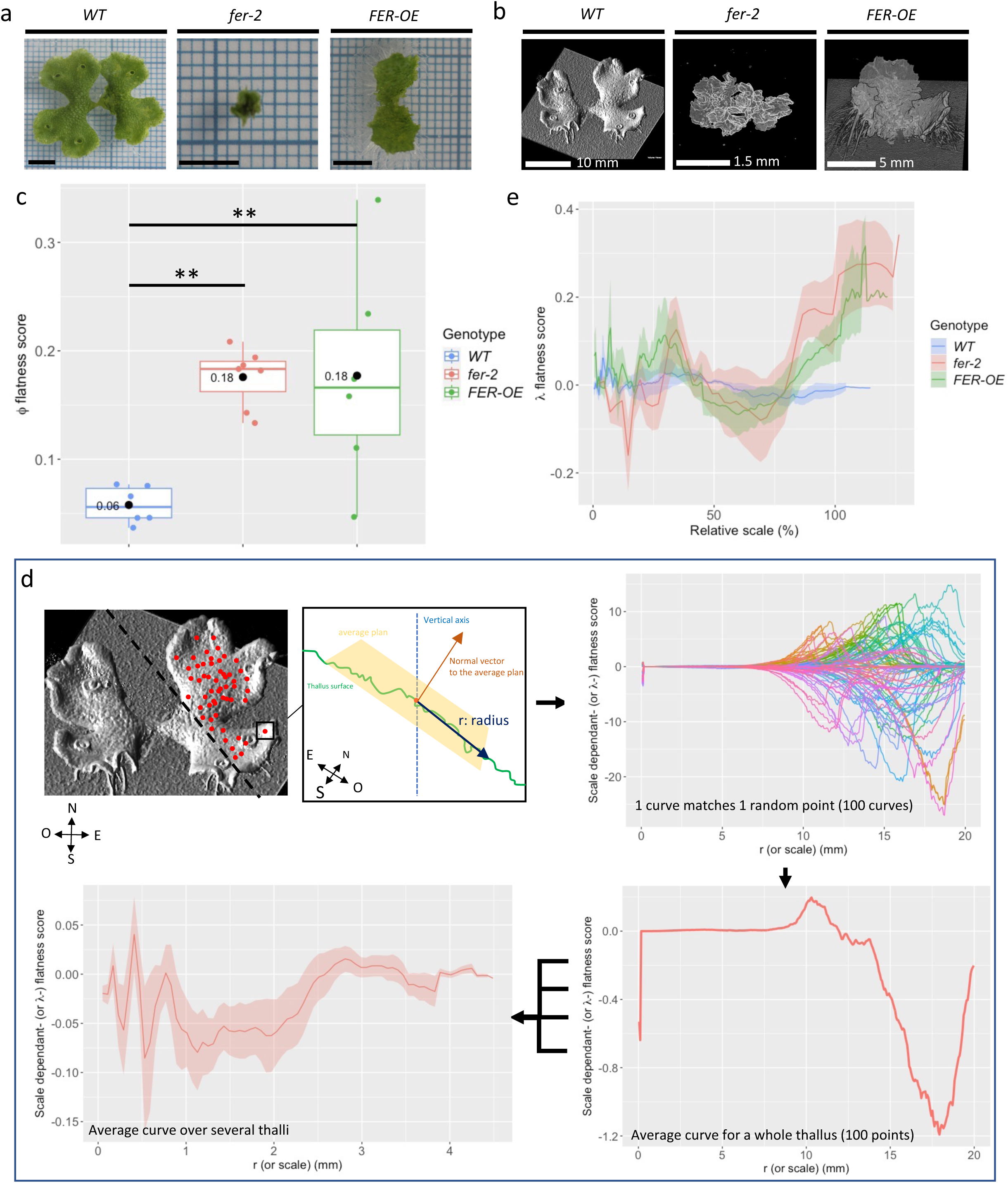
Flatness of Marchantia thalli with or without altered expression of *FERONIA*. **a**, **b** and **c**) Pictures and flatness scores of WT, *fer-2* mutant and *FER-OE* transgenic plants; **a**) Pictures of 15-day-old thalli. Scale bar = 5 mm; **b**) µCT scan surface-reconstruction of 16-day-old thalli; **c**) ϕ flatness score of 16-day-old thalli (6 WT, 7 *fer-2* and 6 *FER-OE* thalli). Pairwise Wilcoxon test with Bonferroni correction: **: 0.001 < p-value ≤ 0.01. Black dots indicate the mean value; **d**) Cartoon explaining the approach to the calculation of the λ scale-dependent flatness score (SD-λ flatness score); **e**) SD-λ flatness score of WT, *fer-2* mutant and *FER-OE* transgenic plants, plotted in function of length scale r normalised by the square root of the average thallus projected area for each genotype (the same individuals as those of c were analysed). The shaded area corresponds to the standard error. The absence of a shaded area at high scales indicates that N=1.

To improve our understanding of the alteration of flatness in *fer-2* mutant and transgenic lines, we developed a second metric. The scale dependent flatness score (hereafter referred to as ‘λ flatness score’ or simply ‘λ’) is calculated as the variation of the orientation of the average plane within circular regions of radius ‘r’, for a hundred points randomly selected on the thallus (Eq. **2** and fig **1d**). A hundred curves are thus obtained and then averaged into a single curve characterising the surface of a single thallus. This score is positive for dome-shaped surfaces and negative for saddle-shaped surfaces regardless of the orientation of the curvature and vanishes for flat surfaces (supplementary figure **S1**). The λ flatness score is expressed as a function of the radius ‘r’, thus yielding a curvature across length scales (region sizes). To allow for comparison between genotypes and conditions, ‘r’ is expressed as the percentage of the average thallus size for each genotype and condition (hereafter referred to as the ‘relative scale’). The λ flatness score of multiple thalli can then be combined into an average curve and its standard error (shaded area) characterising flatness at multiple scales for a single genotype and growth condition (fig. **1d** and **e**).

Computation of the λ flatness score confirms that wild type thalli can be considered flat at all scales, with an average λ close to 0. The *fer-2* mutant line displays erratic λ values straying away from that of the wild type and from zero, with noticeably negative and positive values at relative scales of about 12% and 70 to 100 %, respectively, indicating saddle shapes at relatively small scales and dome shapes at larger scales. Similarly, the *FER-OE* transgenic line displays a positive λ score at about 30% and above 80 %, indicating dome-shaped values at these two scales, and possibly reflecting the hyponastic phenotype observed in figure **1b** at larger scales (fig. **1d**). Similar behaviours were observed when looking at absolute scale values (supplementary figure **S2**). Altogether, removing or increasing *FER* expression alters flatness, at several scales.

### *FERONIA* regulates thallus growth through mechanical responses

In *fer-2* mutant thalli, rhizoids burst (Mecchia et al., 2022), consistent with the role of FERONIA in cell wall integrity sensing determined in Arabidopsis (Höfter et al., 2015; Feng et al., 2018). In the *fer-2* mutants of Marchantia, turgor pressure (and so mechanical stress) would be high enough to irreversibly damage the cell wall. We hypothesised that the alteration of thallus flatness is also partially due to high mechanical stress. To test this hypothesis, we changed the agar concentration of the growth medium to reduce the mechanical stress exerted on the cell wall. High agar concentration (2.5%) reduces the water availability in the medium, and therefore, mechanical tension on the cell wall, by reducing turgor pressure or by affecting cell geometry (Ghashghaie et al., 1991; Owens and Wozniak, 1991; Verger et al., 2018). Conversely, we also tested the effect of higher mechanical tension in the cell wall by using a low agar medium (0.6%) (Ghashghaie et al., 1991; Owens and Wozniak, 1991; Verger et al., 2018).

Consistently with an increased (reduced) cell wall tension, *fer-2* thalli are larger (smaller) when grown on low (high) agar medium (fig. **2a** and **c**), in agreement with the trends in thallus volumes and weight across conditions (supplementary figure **S3**). However, the *FER-OE* transgenic line and the WT line show no significant changes in their overall size, suggesting that they compensate for the changes in cell wall tension induced by changes in agar concentration. In particular, this implies that the *FER*-*OE* transgenic line conserved the capacity to adapt to changes in the mechanical status of the plants.

**Figure 2:**
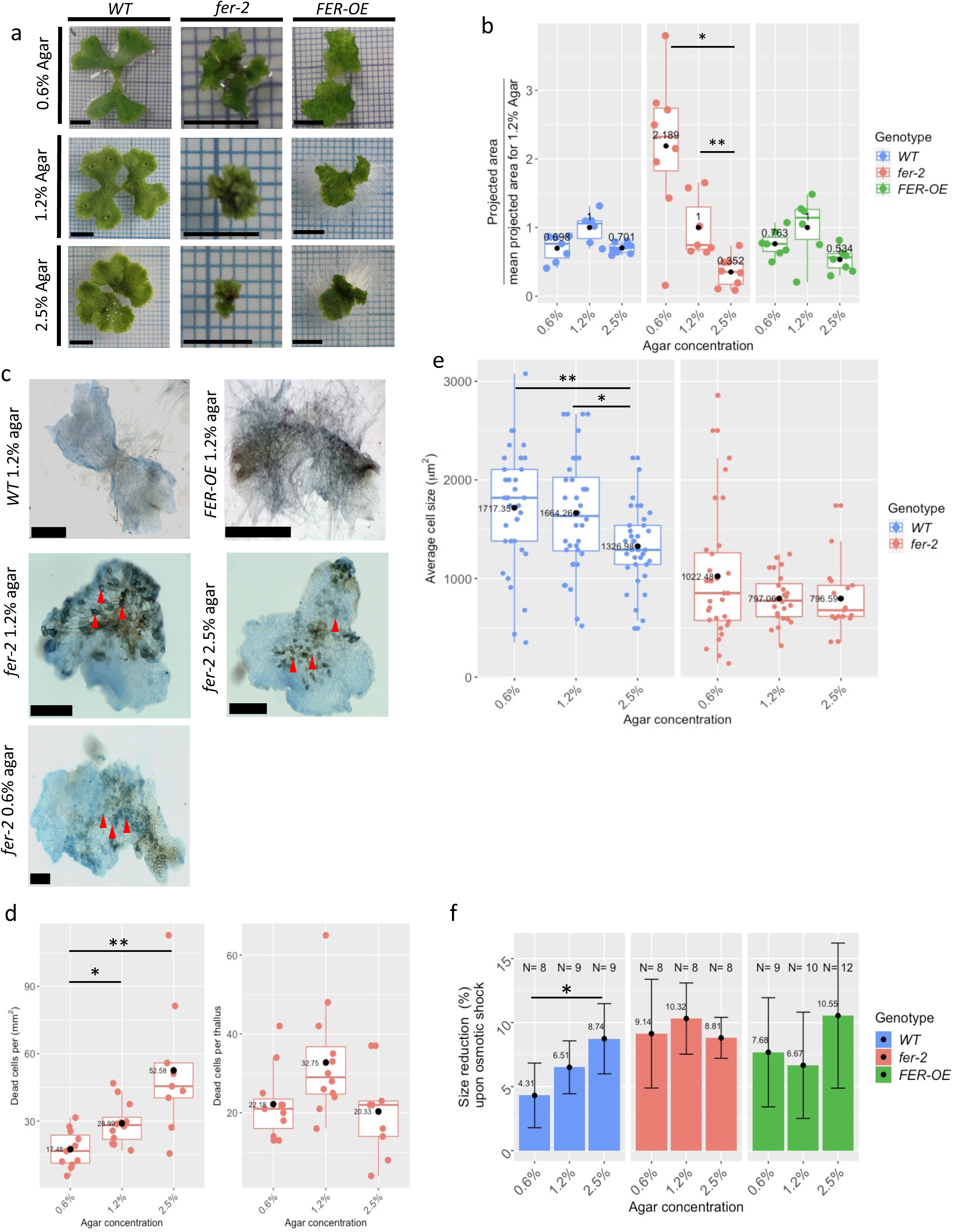
Effect of changes in agar concentration of culture medium on size, and mechanics of thalli. Data for WT, *fer-2*mutant and *FER-OE* transgenic plants grown on 0.6%, 1.2% (standard concentration) and 2.5% agar media. a) Macroscopic pictures. Scale Bar = 5 mm; b, e, d and f) Black dots indicate the mean; b) Measurement of the projected area of 16-day-old thalli: 6 WT, 7 *fer-2* and 6 *FER-OE* thalli grown on 1.2% agar medium, 8 WT, 8 *fer-2* and 8 *FER-OE* thalli grown on 2.5 % agar medium, and 7 WT, 8 *fer-2* and 7 *FER-OE* thalli grown on 0.6 % agar medium. A pairwise Wilcoxon test with Bonferroni correction was used: *: 0.01 < p-value ≤ 0.05; **: 0.001 < p-value ≤ 0.01; c) Trypan blue coloration of 8-day-old *fer-2* thalli grown on low, standard, and high agar concentrations –dark blue coloration indicates dead cells (red arrow heads). Trypan blue coloration exclusion test performed on wild type thalli shows no visible cell death on standard agar concentration. Scale bar = 250 μm; d) Number of dead cells per mm^2^ in 8-day-old thalli (left), and number of dead cells per thallus measured in the same individuals (right). Pairwise Wilcoxon tests with Bonferroni correction were performed: *: 0.01 < p-value ≤ 0.05; **: 0.001 < p-value ≤ 0.01; e) Cell size measured in 14-day-old thalli; f) Size reduction in 7-day-old thalli after an osmotic shift in 250mM mannitol for 10mn – larger size reduction indicates lower cell wall stiffness or higher turgor pressure. A pairwise Wilcoxon test was performed with a Bonferroni correction: *: 0.01 < p-value ≤ 0.05.

We next examined the cellular basis of changes in plant size upon changes in agar concentration. It was shown that *fer-2* mutants are characterised by the presence of dead cells (Mecchia et al., 2022). We therefore sought whether changes in *fer-2* thallus size could be ascribed to cell death. We performed a trypan blue exclusion test on 8-day-old mutant thalli to quantify cell death. We observed no cell death in wild type (fig **2c**). In *fer-2*, the number of dead cells per unit area is three times lower on low agar medium than on high agar medium (fig. **2d**) (17.48 dead cells/mm^2^ on 0.6% agar against 52.58 dead cells/mm^2^ on 2.5% agar medium). This would suggest that the increase in thallus size is due to less cell death. However, cell death occurs during the early developmental phases (Mecchia et al., 2022) and it could be that a low number of dead cells per unit area is the result of a similar number of dead cells spread over larger surfaces. To test this hypothesis, the total number of dead cells was counted in the same individuals. No significant difference can be observed (fig. **2d**). Moreover, the number of dead cells per mm^2^ (mostly in the range 10-50) is much smaller than the number of epidermal cells per mm^2^ (approximately 1250 cells/mm^2^ for *fer-2* grown on 2.5% agar medium and 1000 cells/mm^2^ for *fer-2* grown on 0.6% agar medium, estimated from the data shown hereafter), which cannot directly account for 2-fold changes in thallus size. This suggests that cell death does not explain changes of *fer-2* thallus size in response to changes in agar concentration.

Cell size was then assessed by counting the number of cells within defined surface area between 100 × 100 μm^2^ and 200 x 200 μm^2^, thallus size permitting. In wild-type, cell size decreased with agar concentration, like thallus size. Interestingly, *fer-2* mutants grown on high or low agar media do not show any significant differences in cell size (Fig **2e**), indicating that the larger *fer-2* thalli observed on low agar medium do not result from larger cells.

The absence of variations in the thallus size and cell size of *WT* thalli grown on high or low agar media suggests that the mechanical properties of the cell wall adapt to variations in turgor pressure to maintain cell growth and cell size. Given the dramatic changes of thallus size in *fer-2* grown on low agar medium, and the lack of variation in cell size between *fer-2* grown on high and low agar, we sought whether *fer-2* mutants retained some capacity to adjust their cell wall mechanical properties to changes in turgor pressure. Interestingly, cellular force microscopy revealed lower stiffness for *fer-2* thalli, indicating that cell wall stiffness and/or turgor pressure are lower in this mutant (Mecchia et al. 2022). We therefore hypothesised that FERONIA is required to adjust cell wall stiffness or turgor to maintain growth and flatness when cell wall tension is altered.

To test this hypothesis, we grew thalli on high, standard or low agar concentrations for 7 days, and quantified the cell wall elastic properties by measuring size reduction of the thallus upon a shift from normal isotonic solution to a hypertonic solution (isotonic solution supplemented with 250 mM mannitol). Softer cell walls will result in higher size reduction. WT thalli show size reduction that scales with the agar concentration (4.31% on low agar and 8.74% on high agar) fig. **2f**, indicating that the cell wall stiffness is higher on low agar medium than on high agar medium. This is consistent with an adjustment of tissue mechanics to external osmolarity. The *fer-2* mutants showed a complete absence of shrinkage variations across all agar concentrations, showing that *fer-2* fails to adapt the mechanical properties of its cell wall to variations in external osmolarity. Furthermore, size reduction observed in *fer-2* thalli is very close to that observed in WT thalli grown on high agar medium, indicating that *fer-2* has abnormally soft cell wall, consistent with the low cell stiffness measured in Mecchia et al., 2022. The *FER-OE* transgenic line shows no significant difference in size reduction when grown on low or high agar medium, although higher variability between samples makes it more difficult to conclude. This could be consistent with a saturated response of *FER-OE* to mechanical tension. Together, these results suggest that FERONIA is involved in the sensing of mechanical stress and triggers mechanical adaptations, leading to an adjustment of thallus size independently of cell size or cell death.

### Specific regulation of *FERONIA* is required for the production of flat thalli

To test the impact of cell wall tension in the maintenance of flat shape, we further characterised thallus flatness in plants grown on 0.6%, 1.2% or 2.5% agar medium. Calculation of the ϕ score does not indicate significant changes of the flatness of in WT thalli (fig. **3a**), confirming adaptation to changes in cell wall tension. The *fer-2* mutant also displays no variation in flatness ϕ score. However, further examination of the λ flatness score indicates flatter surfaces for *fer-2* thalli grown on high agar and less flat (curvier) surfaces for thalli grown on low agar medium (fig. **3c** and supplementary figure **S4**). This suggests that, in the absence of *FER* function, higher cell wall tension induces loss of flatness, consistent with the hypothesis that buckling is induced by mechanical stress. *FER-OE* thalli show improved ϕ flatness score on high agar medium: 0.1 on 2.5% agar compared to 0.18 on 1.2% agar medium (fig. **3a**), as well as lower λ flatness score at all scales (fig. **3d** and supplementary figure **S4**), suggesting that *FER-OE* plants lose flatness by over-responding to high cell wall tension. Altogether, these results suggest that intermediate levels of FERONIA are required to adjust growth rates and to prevent buckling by responding to mechanical stress.

**Figure 3:**
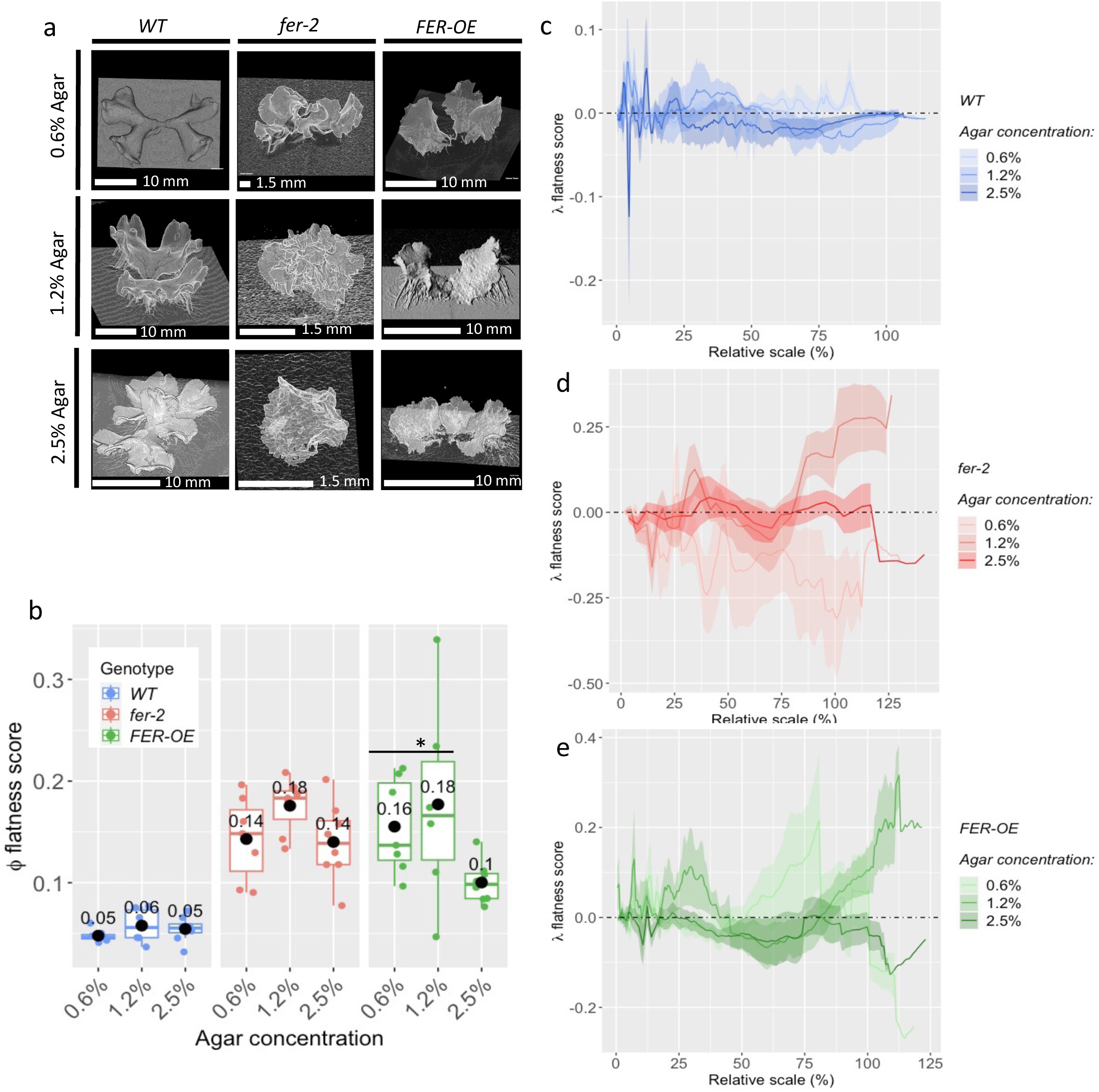
Flatness in low, standard and high agar concentrations: **a**) µCT scan surface-reconstruction of 16-day old thalli; **b**-**e**) The same individuals were used to calculate the ϕ and λ scores as those of figure 2c: 6 WT, 7 *fer-2* and 6 *FER-OE* thalli grown on 1.2% agar medium, 8 WT, 8 *fer-2* and 8 *FER-OE* thalli grown on 2.5 % agar medium, and 7 WT, 8 *fer-2* and 7 *FER-OE* thalli grown on 0.6 % agar medium. **b**) ϕ flatness score of 16-day-old thalli. A pairwise Wilcoxon test with Bonferroni correction was used: *: 0.01 < p-value ≤ 0.05. Black dots indicate the mean value; **c**, **d,** and **e)** λ flatness score plots in function of the relative scale for each growth condition and genotype. Shaded area represents the standard error. The absence of shaded area for high relative scales indicates that N=1; **c**) WT; **d**) *fer-2*, **e**) *FER-OE*.

### Application of external mechanical pressure partially restores thallus size in *fer-2* mutants

Given that an increase in agar concentration made *fer-2* and *FER-OE* plants slightly flatter, we wondered whether direct reduction of cell wall mechanical stress would restore flatness of *fer-2* and *FER-OE* thalli. We hypothesised that compressing thalli with a film would reduce the mechanical stress exerted on the cell wall by helping the cell wall to resist turgor pressure at the contact points between the film and the thallus (fig. **4a**). Transparent thin films of polydimethylsiloxane (PDMS) were selected to allow for gas exchange (Adler et al., 2010) and reduce other sources of abiotic stress.

**Figure 4:**
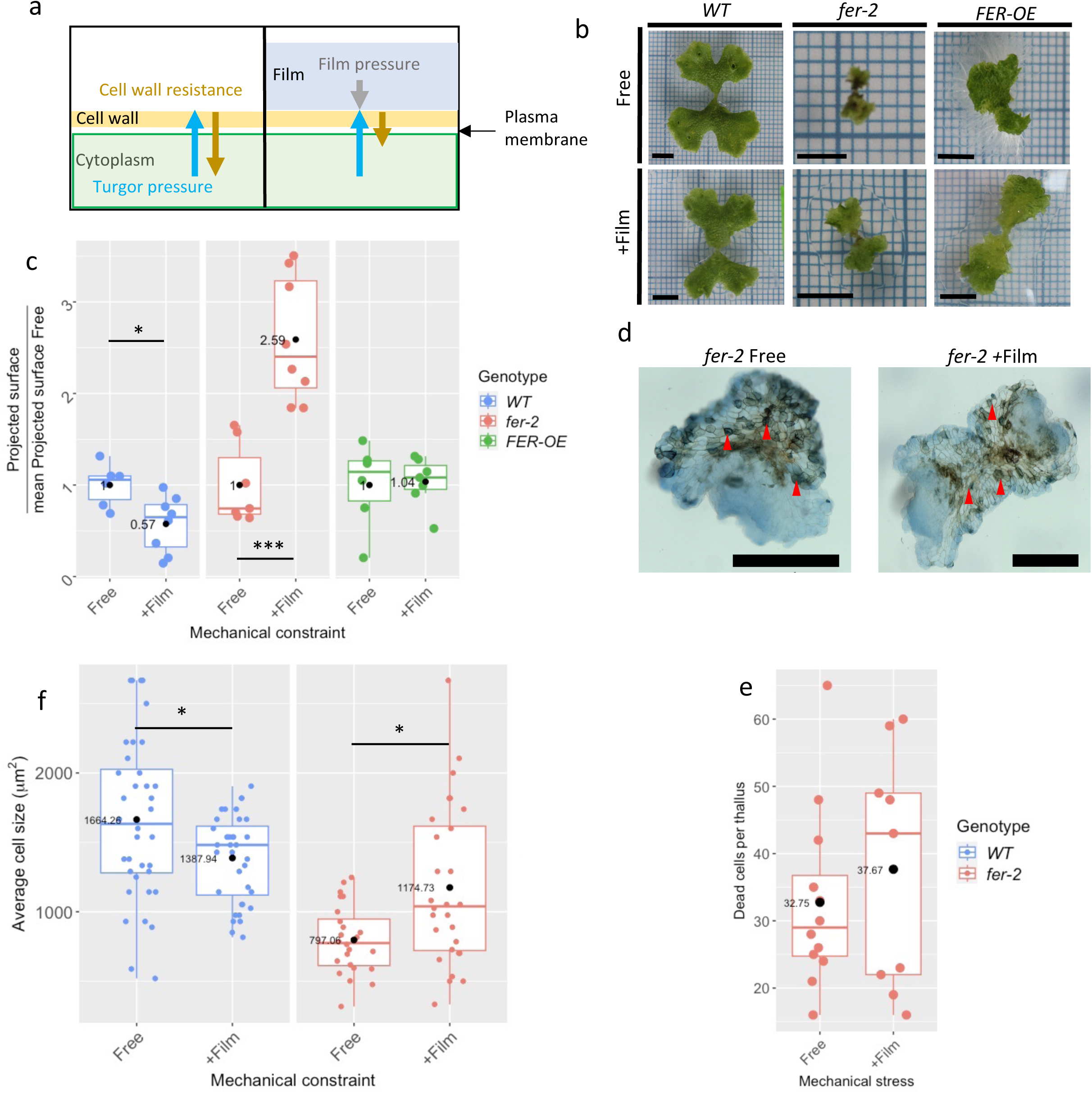
Effect of thallus compression by PDMS film on thallus size, cell size, and cell death. **a**) Cartoon showing how a topping PDMS film can contribute to the reduction of mechanical stress on the cell wall. Blue arrows show the turgor pressure exerted on the cell wall. Brown arrows show the cell wall resistance to the inner turgor pressure. Grey arrow shows the mechanical pressure exerted by the PDMS film on the cell wall at contact points; **b-f)** Qualitative and quantitative analyses of plants grown with (+Film) and without film (Free). **b**) Macroscopic pictures of 15-day-old thalli; **c**, **d,** and **f)** Black dots indicate the mean value. **c)** Projected surface of 16-day-old thalli: 6 WT, 7 *fer-2* and 6 *FER-OE* thalli grown without film, 8 WT, 8 *fer-2* and 7 *FER-OE* thalli grown with film. WT: a t-test was performed: *: 0.01 < p-value ≤ 0.05; *fer-2*: a t-test was performed: ***: p-value ≤ 0.0001, *FER-OE* a Wilcoxon test was performed: *: 0.01 < p-value ≤ 0.05; **d**) Trypan blue coloration exclusion test performed on 8-day-old *fer-2* thalli grown on low, standard, and high agar concentrations – dark blue coloration indicates dead cells (red arrow heads). Scale bar = 500 μm; **e**) Number of dead cells counted in 8-day-old thalli; **f**) Cell size measured in 14-day-old thalli. A t-test was performed to compare WT samples and a Wilcoxon test was used to compare *fer-2* mutant samples *: 0.01 < p-value ≤ 0.05.

Measurement of the thallus surface in the wild type shows that the application of thin PDMS films significantly reduces the projected surface (about 57% of the projected surface of thalli grown without film) (fig. **4b** and **4c**). Significant reductions in weight and volume are also visible upon application of PDMS films (supplementary fig. **S5**). Surprisingly in the case of the *fer-2* mutant, the projected surface ratio, the volume, and the weight are at least twice as large with films (+Film) than without film (Free) (fig. **4b** and **c**, and supplementary fig. **S5).** No significant difference was observed in the case of *FER-OE* size with any metrics. Altogether, this shows that the application of thin film restores thallus growth in *fer-2* mutants (fig. **4d** and supplementary fig. **S5).**

As for the variations in thallus sizes observed on various agar concentrations, we sought the cellular basis of the changes in thallus shape grown with and without PDMS films. We wondered whether this involves cell death, or cell expansion. The number of dead cells per thallus is not significantly affected by film application in *fer-2* mutants. This suggests that cell death is limited to the early development of the thallus, consistent with the conclusions from low and high agar experiments. Moreover, as explained earlier, the number of dead cells appears too small when compared to the estimated number of cells per thallus to explain such dramatic changes in thallus size. We then examined cell size in these experiments. Wild type thalli did show a significant decrease of the overall cell size when topped with a film (about 1664 μm^2^ without film, and 1388 μm^2^ with film), whereas the *fer-2* mutant thalli shows larger cells on average when topped with a film (about 797 μm^2^ without film, and 1174 μm^2^ with film) (fig. **4f**). This suggests that application of film partially restores cell size in the mutant. Taken together, these results indicate that the increase in cell size in the *fer-2* mutant topped with a film is the main reason for the improvement of the size the thallus shown in figure 4.

### Application of external mechanical pressure partially restores flatness in *fer-2* mutant and *FER-OE* thalli

We then questioned whether application of external mechanical pressure could result in changes in thallus flatness. Wild-type thalli do not display any significant alteration of flatness and remain equally flat regardless of the presence of a film (fig. **5a**) and their average ϕ flatness score remains unchanged (0.06) (fig. **5b**). A slight change is noticeable regarding their λ curvatures for relative r values above 30%, where the natural curvature is altered by the film, but remains close to zero (fig. **5c** and supplementary figure S6). The *FER-OE* thalli display very little hyponasty upon the application of the film (fig. **5a**). Furthermore, their ϕ flatness scores indicate that the application of the film is sufficient to restore flatness down to the values of the wild type (0.18 without film and 0.06 with film) (fig. **5b**). The λ flatness score indicates a complete loss of curvature at scales above 50 %, which is consistent with the recovery of a normal hyponastic behaviour (fig. **5e** and supplementary figure S6), and a reduction in curvature around a scale of 30%. Altogether, this suggests that over-reaction to mechanical stress in *FER-OE* is tempered by a reduction in wall tension. The *fer-2* mutant also appears flatter upon film application (fig. **5a**). Calculation of the ϕ score also confirms the restoration of flatness (0.18 without film and 0.08 with a film) (fig. **5d**). The application of film decreases the λ curvature at scales around 30 % and restores flatness for scales above 60 % (fig. **5d** and supplementary figure **S6**). Altogether, these results indicate that the application of supplementary mechanical constraints, in the form of a topping thin film, as a way to reduce cell wall mechanical stress through partial compensation of the turgor pressure, succeeds in improving the overall flatness of the thalli in both mutant and transgenic lines.

**Figure 5:**
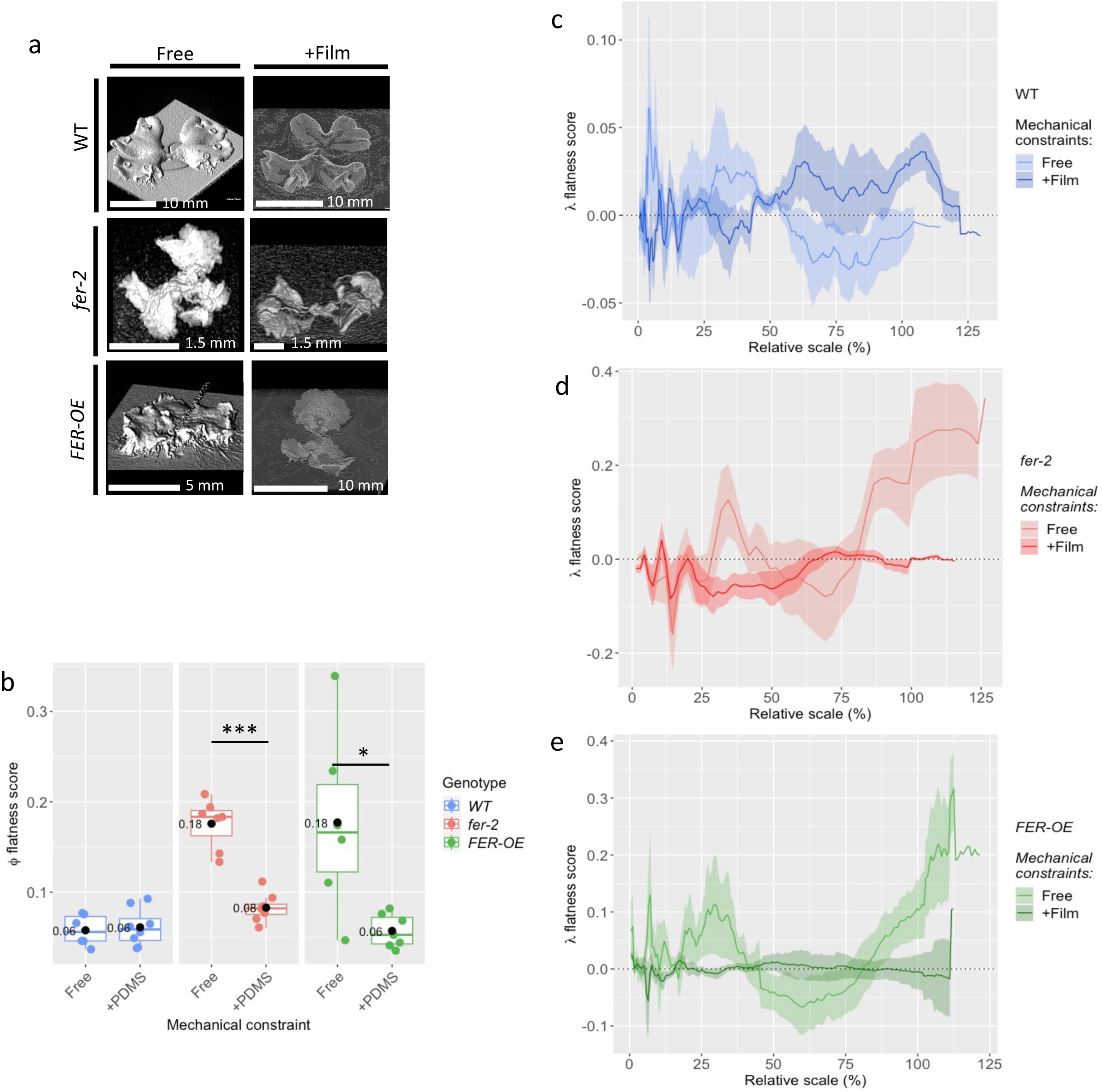
Effect of thallus compression on flatness. **a**) µCT scan surface reconstruction of 16-day-old thalli; **c**-**e**) Quantification of flatness in thalli grown with (+ Film) and without (Free) a topping PDMS film. The same 16-day-old thalli were analysed as those of fig 4d: 6 WT, 7 *fer-2* and 6 *FER-OE* thalli grown without film, 8 WT, 8 *fer-2* and 7 *FER-OE* thalli grown with film; **b**) ϕ flatness score of 16-day-old thalli. *fer-2*: a t-tests was performed: ***: p-value ≤ 0.0001. *FER-OE*: a Wilcoxon-test was performed: *: 0.01 ≤ p-value ≤ 0.05. Black dots indicate the mean value; **c, d** and **e)** Plot of the λ flatness score in function of the relative scale for each growth condition and genotype. Shaded area represents the standard error. The absence of shaded area for high relative scales indicates that N=1; **c**) WT; **d**) *fer-2*, **e**) *FER-OE*.

## Discussion

In this article we bring understanding in the possible ways by which mechanical responses influence organ size and shape. We specifically considered the development of flat organs and developed a new way to quantify flatness (ϕ and λ flatness scores) of complex surfaces. These two metrics helped us to analyse and compare the surface of multiple genotypes and developmental conditions of *Marchantia polymorpha.* We demonstrate that the *feronia-2* mutant and the *FER-OE* transgenic lines are altered in the development of flat thalli and display impaired regulation of the mechanical properties of the cell wall upon variation of cell wall tension. Finally, we show that application of external mechanical constraints using PDMS films, which we expect to reduce mechanical stress exerted by turgor pressure on the cell walls, results in larger cells and restores flatness in the *fer-2* mutants.

In this study, the ϕ score allowed for quick characterisation of thallus surfaces and revealed consistent alteration of flatness in *fer-2* mutant and *FER-OE* transgenic thalli. This metric provides an interesting approach to assess overall surface flatness because of its simplicity and its short computation time (typically less than 20 seconds per sample). Furthermore, it also provides 2-D surface resolution when typical measurement of flatness focuses on unidirectional curvature analysis (Liu et al., 2010; Ren et al., 2018; Natarajan et al., 2020), making the ϕ score an efficient choice when tackling complex surfaces.

The development of the λ flatness score complemented this approach by providing scale resolution. Here, this score has provided fine resolution of thallus curvature and flatness and showed differences in flatness and curvature between *fer-2* mutant thalli grown on 2.5% and 1.2 % agar. This score also provides supplementary characterisation of the curvature and can discriminate saddle-shaped surfaces from dome-shaped surfaces. The Gaussian curvature is also able to discriminate between these two types of surfaces but does not directly provide scores relative to scale. The power of the λ score rests in the fact that it calculates curvature at all scales in a single run and thus gives a single curve characterising the whole surface. Computation time remains however longer than that of the ϕ score (typically about 10-20 min per sample). Together, these scores provide a fine resolution description of the thallus surface and allow for the analysis of broad and subtle changes in the surface aspect of *Marchantia polymorpha* thalli.

The alteration of flatness observed in the mutant and transgenic lines of *FERONIA* shows that intermediate levels of *FERONIA* expression are required for maintenance of flatness. Since *FERONIA* has been identified as involved the maintenance of cell wall integrity, possibly sensing mechanical stress (Mecchia et al., 2024; Xu et al., 2024; ling et al., 2018; Tang 2018, Shi et al., 2014), we suggest that both mechanosensing and cell wall integrity sensing are required for the maintenance and development of proper organ shape.

We showed here that modification of agar concentration is sufficient to trigger modifications in cell wall stiffness in wild type thalli, and that such capacity is lost in *fer-2* mutant and *FER-OE* transgenic lines. This indicates that FERONIA is involved in perception and or response to cell wall tension. This might occur either through the binding of FERONIA to pectin (Lin et al., 2022) or by the binding of RALF-Ligand peptides signalling cell wall damage (Liu et al., 2023). Measurement of tissue mechanics in *fer-2* mutant thalli using cellular force microscopy showed that either Young’s moduli of the epidermal cell wall or turgor pressure are reduced in the *fer-2* mutant (Mecchia et al., 2024). Here, we find that *fer-2* is specifically affected in its cell wall modulus. This supports the hypothesis that FERONIA is necessary to maintain cell wall stiffness and cell wall integrity.

Loss of flatness often is the result of buckling, usually caused by deregulated growth and resulting tissular compressive stress (Sharon et al., 2004, Boudaoud et al., 2010). This suggests that the lack of flatness observed in *fer-2* mutant and *FER-OE* transgenic lines could be the result of abnormal tissue growth, consistent with the abnormal cell size observed in the *fer-2* mutants both in this study and in Mecchia et al., 2022. We also found that treatments that reduce cell wall tension and tissular stress make *fer-2* and *FER-OE* flatter, supporting the hypothesis that loss of flatness is induced by buckling in these genotypes. Furthermore, it was shown that brassinosteroids induce accumulation of FERONIA at the plasma membrane, which results in a feedback loop attenuating brassinosteroid promoted cell elongation (Wolf et al., 2012; Ajeet et al., 2023) thus showing how FERONIA can interact with cell expansion mechanisms. Together, this suggests that *FERONIA* is important for the regulation of cell size and cell expansion.

Interestingly, none of the treatments applied on the *fer-2* mutant thalli successfully prevented cell death, despite some restoration in flatness and size. This suggests that cell death is not detrimental to the acquisition of organ shape and that cell death is not responsible for tissue buckling in this case. However, the cause of cell death remains unclear. The softer cell wall observed in the *fer-2* mutant, reported in both Mecchia et al., 2022 and in this study suggests that cell death is the result of cell burst. Given that cell burst was reported in the rhizoid cell (Mecchia et al., 2022), it is plausible that cell death mostly occurs in rhizoid precursors, consistent with the position of the rhizoid precursor cells and the relatively constant number of dead cells across *fer-2* mutants grown on various media and with or without films. Other factors than the mechanical yielding of the cell wall may also induce cell death, but they have not been investigated here. FERONIA is often reported as playing a role in plant immune response (Gao et al., 2018, Jing et al., 2022, Yang et al., 2020), suggesting that misregulation of *FERONIA* expression could cause abnormal immune response resulting in cell death. The tight regulation of cell expansion and proliferation is crucial to the maintenance of organ size and shape (Powell and Lenhard 2012). This suggests that the recovery of normal cell size in the *fer-2* mutants topped with films contributes to the restoration of normal thallus size. Interestingly, *FERONIA* was found to play a role in determining cell size by preventing cell wall acidification and attenuating brassinosteroid-induced cell expansion in *Arabidopsis thaliana*, possibly to prevent cell burst (Chaudhary et al., 2023). Here, we have discussed flatness in organs such as sepals, leaves, and thalli, all of which have constant thickness and are relatively thin. A prerequisite is thus the initiation of thin organs, which may depend on oriented cell divisions and/or on organisation of cortical microtubules (Zhao et al., 2020; Wallner et al., 2024). Furthermore, changes in thickness have been correlated with alterations of flatness in the sepals of the *asymmetric leaves 2 (as2)* mutants (Yadav et al., 2023) suggesting that flatness requires constant and small thickness. The importance of coordinated growth between the adaxial and abaxial layers in *Arabidopsis* sepals (Yadav et al., 2023) also suggests that alteration of flatness in the *fer-2* mutant and *FER-OE* transgenic lines could result from lack of mechanical feedback between cell layers thus potentially promoting buckling.

In the *angustifolia* mutants of Marchantia (Furuya et al., 2018), lack of thallus flatness was associated with abnormal orientation of the cortical microtubules at the apical notches. This suggests that microtubule orientation is involved in the maintenance of thallus flatness and raises questions on the possible mechanisms triggered by MpFERONIA in the maintenance of flatness. However, single cell laser ablations of epidermal cells of the Arabidopsis *fer-4* mutants showed normal reorientation of cortical microtubules in the neighbouring cells, suggesting a *FERONIA* independent mechanism for the regulation of microtubule arrangements (Malivert et al., 2022).

Here we have focused on the mechanical regulation of flatness. It was suggested that A*tKTN1* (Uyttewall et al., 2012) and the *VIP3* (Jensen et al., 2016; Trinh et al., 2024) mutants are involved in the robustness of organ shape development via the promotion of cell growth and proliferation heterogeneity. Given their involvement in the mechanical perception and response to mechanical stimuli, and the role of *FERONIA* in mechanical stress perception, we raise the question of a possible role of *FERONIA* in the robustness of flat organ developments via the maintenance of cell growth heterogeneity. Future work should also reveal whether and how such mechanical regulation operates with pathways that coordinate growth between neighbouring cells and across an organ, possibly ensuring the robustness of morphogenesis of flat organs.

## Material and Methods

### Plant material and growth conditions

The Tak-1 plant was used as wild-type (WT) and propagated by cutting onto 1.2 % agar (Duchefa Biochemie) solid medium containing 0.2% Gamborg B5 supplemented with vitamins (Duchefa Biochemie) and incubated under constant light (around 40 μmol.photons.m^−2^.s^−1^). The *fer-2* mutant line as well as the _pro_*MpEF1::MpFERONIA-mCitrine* transgenic line 9 (*FER-OE*) were provided by Martin A. Mecchia and were characterised in (Mecchia et al., 2022). Mutant and transgenic lines of *FERONIA* were maintained by cutting and grown under the same conditions as WT, except that the medium was supplemented with 1% sucrose (Sigma Aldrich) to boost the production of gemmae.

### Imaging techniques

Gemmae were photographed at day 15 using the close-up standard setting of a Canon® camera and imaged by X-ray micro computed tomography (μCT) at day 16 using the Quantum FX, Perkin Elmer Micro-CT (90 kVp (kiloVolt peak) and 160 mA current for 3 minutes). Various μCT field views were used depending on the overall size of the thallus: width of 5, 10, 20 and 30 mm for a resolution of 10, 20, 40 and 60 μm, respectively. Optimal field view was used for each sample. μCT scan surface reconstruction was performed using the ‘volume viewer’ plugin of ImageJ on raw .tiff stacks of images obtained from Micro-CT scans.

### Surface detection and calculation of the flatness scores

Stacks of images were binarised and resliced along the z axis before loading into MorphoGraphX (https://morphographx.org/). Voxel size was manually corrected to fit MicroCT scan resolution and a ‘Gaussian blur stack’ with σ comprised between 5 and 30 μm was implemented before surface detection using the ‘Edge detect’ action. A mesh was then created by triangulation using the ‘Marching cube surface’ item with a knot-distance of 60 μm. x,y and z coordinates were then exported in ‘.txt’ format and pre-processed using RStudio (https://posit.co/). Pre-processes include the correction of the artificial possible tilting of the thallus using linear regression and filtering for the maximal z-value for each xy coordinate to avoid artificial walling induced by the surface extraction in MorphographX.

The standard deviation of the Z-coordinates σ was calculated using RStudio after surface extraction and pre-processing. Projected area was calculated using ImageJ (https://imagej.net) on Z-projected stacks. *φ* score was calculated as in Eq. (1).

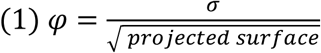

The λ flatness-score assesses flatness at the given scale ‘r’ and is computed as an average over disks of size r. It was calculated for a 100 points selected randomly using Python and is derived from the x,y and z-coordinates of the normal vector (components *nx*, *ny* and *nz*) to the average plane approximating the thallus surface calculated for each disk of radius r centred on one of the 100 random points. The ‘r’ radius was incremented by 30 μm until reaching the square root of the projected area (2). The λ curvature score was defined as in Eq. (2)

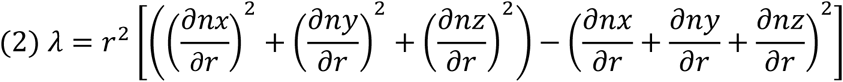

and was calculated using Python. One ‘λ’ curve was obtained for each of the 100 points of a surface. The curves were then averaged together for each value of ‘r’. Finally, The average λ flatness score was once more averaged over all the individuals sharing similar genotypes and growth conditions. The standard error was then calculated as in Eq. (3) with ‘N’ the number of individuals and σ_λ_ the standard deviation of λ, to account for variability among populations.

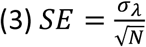

### Cell death assays

Trypan blue exclusion test was used to mark dead cells in 8-day-old gemmae as performed in (Mecchia et al., 2022), with an incubation time in trypan blue dye between 15 and 30 seconds to reduce background noise.

### Mechanical constraints assay

To apply mechanical constraints on thalli, a thin film of Poly(dimethylsiloxane) (PDMS) was placed onto 4-day-old gemmae. PDMS films were produced by mixing 12 g of SYLGARD™ 184 Silicone Elastomer Base (or PDMS) and 4g of SYLGARD™ 184 Silicone Elastomer curing Agent (3:1 ratio). The PDMS and curing agent were then vigorously mixed together and placed in a vacuum bell for about 30 minutes. The PDMS mix was then spread onto twice-coated 3M™ NOVEC™ (1720 Electronic Grade Coating) glass slides by spin coating for 30s at 1500 rpm using a POLOS (by SPS) spin coater. Glass slides were then incubated at 75°C overnight to ensure complete curing of the PDMS. Films of 2.4 x 2.4cm were then cut out and autoclaved between Whatman paper discs.

NOVEC ™ coating was performed by rinsing glass slides with NOVEC ™ solution and heating at 110°C on a hot plate for 10 min to evaporate the solvent. This coating process was repeated twice to ensure minimal adhesion between glass and the PDMS films.

### Thallus weight

Gemmae were grown for 17 days on 0.2% Gamborg B5 medium including vitamins and either 0.6%, 1.2% or 2.5% agar. A PDMS film was placed at day 4 to test the effect of external mechanical constraint on some of the thalli grown on 1.2% agar medium. Thalli were plucked from the medium, and the medium was carefully removed from the rhizoids. A New Class MS Mettler Toledo scale with a precision of 10 μg was used to weigh the plantlets.

### Thallus volume

Thallus volume was extracted from stacks of images obtained using μCT. Stacks were then opened using ImageJ and cropped to fit thallus. Images were blurred using Gaussian blur with a radius of 2μm. Background was then removed using a ball radius between 20 and 60 pixels. Stack was then binarised with automatic threshold calculation. Thallus volume was then extracted using the 3D OC Options of the analyse menu. Select object counter from the analysis menu was then used with the statistics and summary options.

### Osmotic shifts

Thalli were grown for 7 days on 0.2% Gamborg B5 medium including vitamins and either 0.6%, 1.2% or 2.5% agar. Thalli were then collected and mounted between glass slide and round cover slip in an isotonic solution of 0.2% Gamborg B5 medium supplemented with vitamins. Thalli were imaged using the Zeiss Axiozoom V16 with a X 10 zoom. Liquid gamborg solution was then removed by capillarity and replaced by a hypertonic solution of 250mM mannitol (Sigma Aldrich) and 0.2% Gamborg B5 liquid solution supplemented with vitamins and liquid edible red dye (brand: ‘Sainte Lucie’, ingredients: azorubine E122 (1,1%), ponceau red E124 (1%), acidifiers: citric acid E330, preservative: potassium sorbate E202 (0,1%)) (diluted a 100 times) to avoid any confusion between the isotonic and the hypertonic solutions. Thalli were imaged after 10 min in the hypertonic solution with the exact same setting as with the isotonic solution.

### Cell size measurement

Cell size was calculated by counting the number of cells in 200×200μm quadrats and by dividing the quadrat area by the number of cells that were counted. Between 12 and 13 quadrats were sampled per WT thallus and three thalli were selected per condition (for a total of 36 quadrats per condition). Cell size was assessed in a similar way in *fer-2* mutant thalli but smaller thalli occasionally required a smaller quadrat size (127×127μm) to accommodate for the size of the accessible region and the high variation in surface height. Fewer quadrats were also often sufficient to cover the entire thallus. Therefore, a fourth thallus was often added to increase the number of measurements. Hence, cell size was estimated in 5, 7, 12 and 8 quadrats over four thalli for 0.6% agar medium; 7, 4 and 13 quadrats over three thalli for 1.2% agar medium; 4, 4, 12 and 8 quadrats over four thalli for 1.2% agar medium + PDMS film, and 9, 1, 3 and 4 quadrats over four thalli for 2.5% agar medium.

### Statistics

All plots and statistical analyses were performed using RStudio and the ‘ggplot2’ and ‘dplyr’ packages. Box plots indicate quartiles and medians. When present on graphs, black dots were separately computed to indicate the mean. Dot plots overlay indicate Y-values and represent all measurements that were performed.

Statistical tests were performed either with a non-parametric test of Wilcoxon or a parametric Student’s t-test. t-tests were performed after verification of the normal distribution of the samples using a Shapiro-Wilk test, and homoscedasticity using a Fisher F-test. When comparing multiple samples, parametric and non-parametric pairwise tests were used with a Bonferroni correction to reduce the detection of false positives.

## Author contributions

**Jordan Ferria:** Plant maintenance, μCT scans, computation of the ϕ and λ scores and script writing, cell size measurement, cell death assays, projected surface measurements, thallus weighing, PDMS film production, graphic plotting and statistical tests, experimental design and paper writing.

**Carla JA Fournié:** Development of the python script calculating λ flatness score and calculation of λ flatness score. Development and optimisation of protocols for the production of PDMS thin films.

**Magdalena H. Jankowska:** measurement of thallus volume.

**Doron Grossman:** Conceptualisation of the λ flatness score.

**Adrienne HK. Roeder:** Paper reviewing and supervision.

**Stéphanie Drevensek:** Experimental design, paper reviewing and supervision.

**Arezki Boudaoud:** Research funding, experimental design, paper reviewing and supervision.

## Acknowledgments

This work was funded by the ‘Agence Nationale de la Recherche’ (FR), Grant #ANR-21-CE30-0039-01. We would like to thank Lotfi Slimani and Baptist Casel for their technical support during μCT imaging at the ‘Université Paris Cité’ (France), as well as Abdillah Mohamed for his technical support, and Elise Muller for her help in managing plant availability and her suggestions on experimental work.

## Supporting data

The supporting data for this paper are available at **10.5281/zenodo.13981438**. Repository contains μCT scan files, excel tables for volume, thallus projected surface, thallus weight, cell size, cell death, Z-coordinate standard deviation, and the python script used to calculate the λ flatness score.

---

**Supplementary figure S1:**
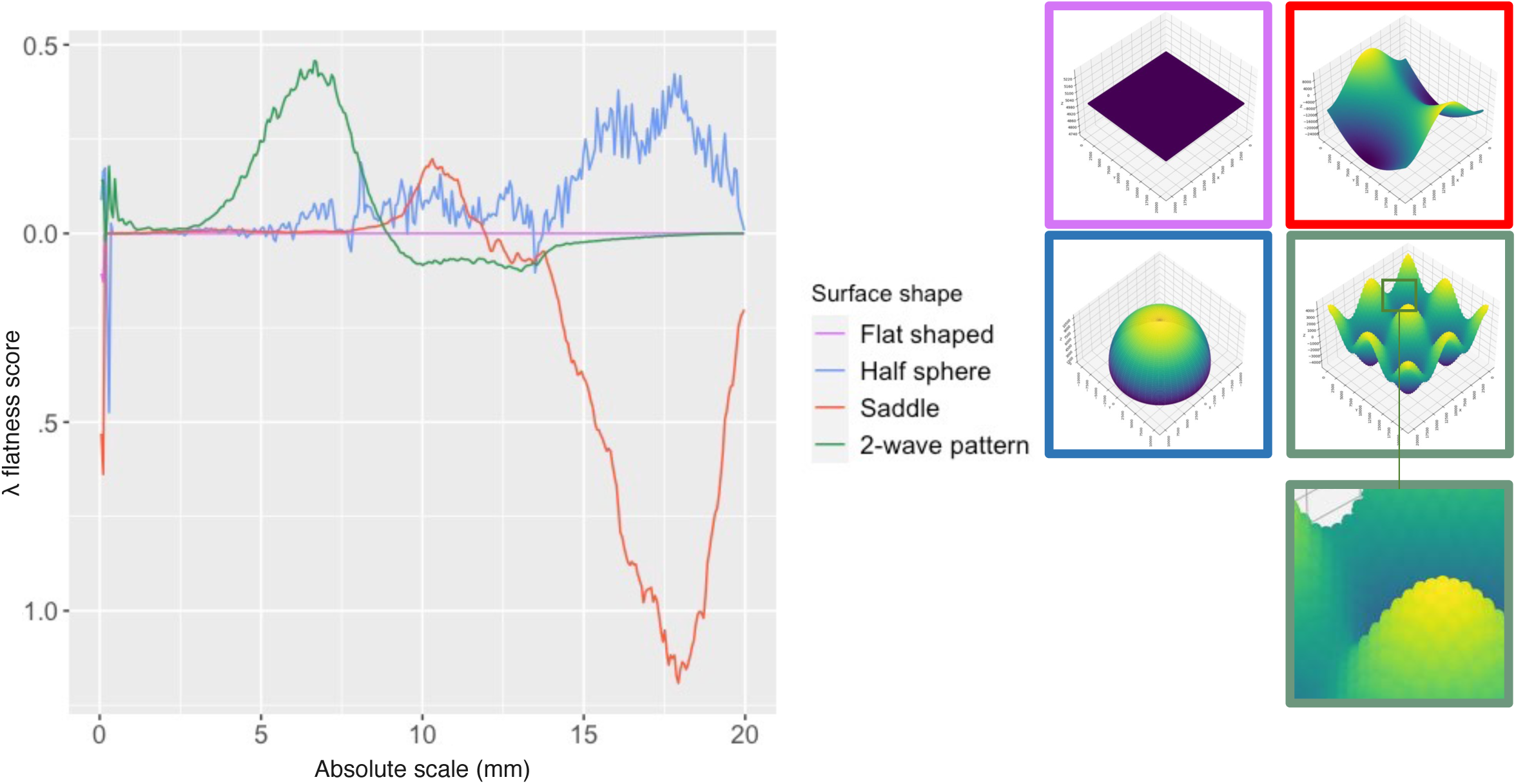
λ flatness score computed on flat, dome shaped, saddle shaped and wavy computed generated surfaces.

**Supplementary figure S2:**
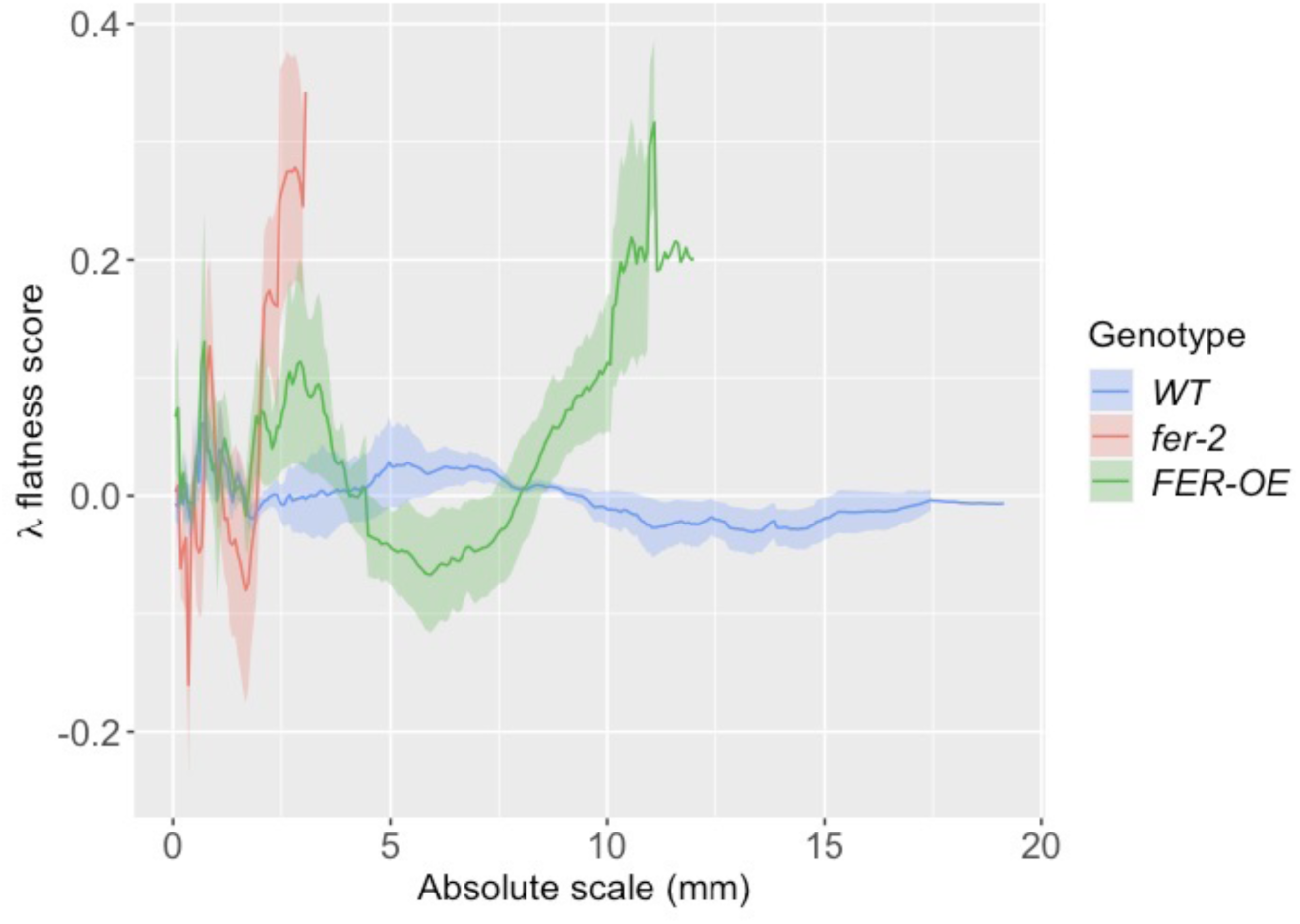
λ flatness score of WT, *fer-2* and *FER-OE* 16-day-old thalli (number of thalli per condition is the same as in figure 1e); The absence of shaded area at high scales indicates that N=1.

**Supplementary figure S3:**
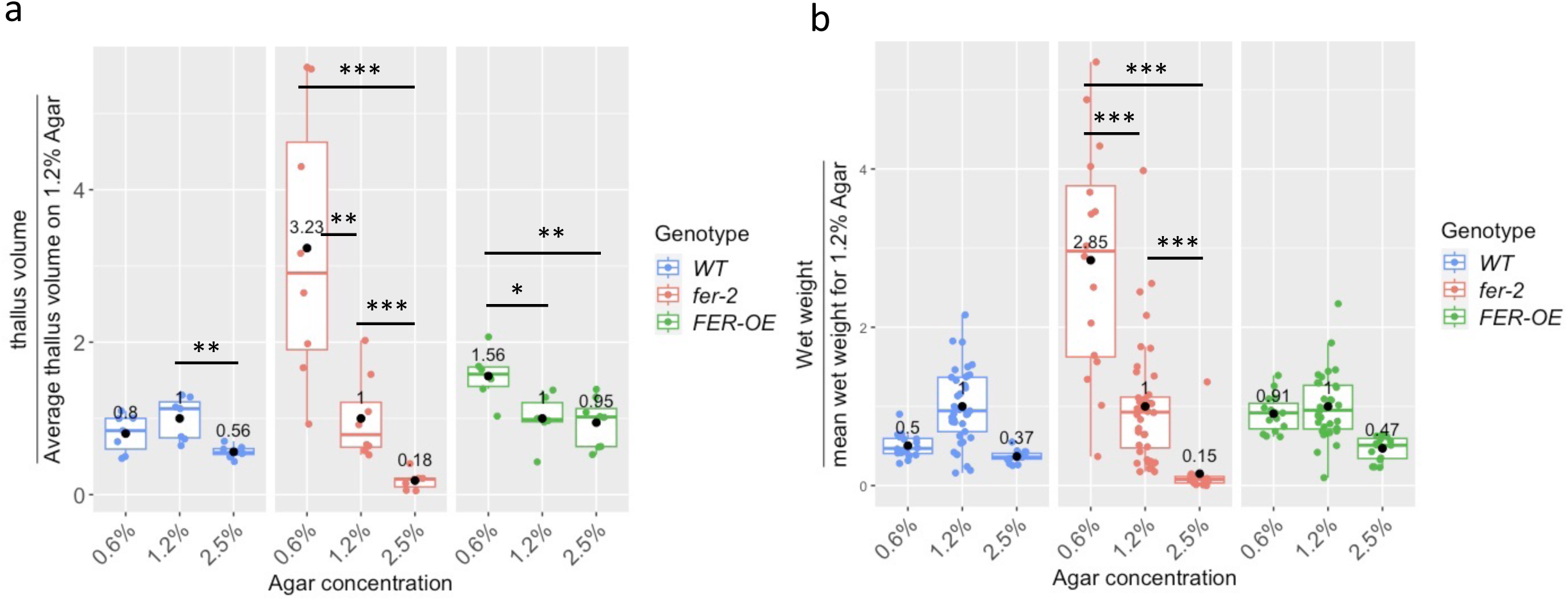
Measurement of thallus weight and volume in thalli grown on 0.6, 1.2 and 2.5% agar medium;**a** and **b**) Black dots indicate the mean.; **a**) Volume of 16-day-old thalli (number of thalli analysed per condition is the same as in figure 2e)(Statistics: WT: pairwise Wilcoxon test with Bonferroni correction was performed **: 0.01 > p-value ≥ 0.001; *fer-2:* pairwise Wilcoxon test with Bonferroni correction was performed: **: 0.01 > p-value ≥ 0.001; ***: 0.0001 > p-value; *FER-OE*: pairwise t-test with Bonferroni correction were performed. *: 0.05 > p-value ≥ 0.01; **: 0.01 > p-value ≥ 0.001; **b**) Weight of 17-day old thalli (pairwise Wilcoxon test was performed with Bonferroni correction ***: 0.0001 > p-value). Number of thalli analysed: (16 WT grown on 0.6% agar medium; 37 WT thalli grown on 1.2% agar medium, 15 WT thalli grown on 0.6% agar medium; 16 *fer-2* grown on 0.6% agar medium; 40 *fer-2* thalli grown on 1.2% agar medium, 15 *fer-2* thalli grown on 0.6% agar medium; 15 *FER-OE* grown on 0.6% agar medium; 32 *FER-OE* thalli grown on 1.2% agar medium, 16 *FER-OE* thalli grown on 0.6% agar medium.

**Supplementary figure 4:**
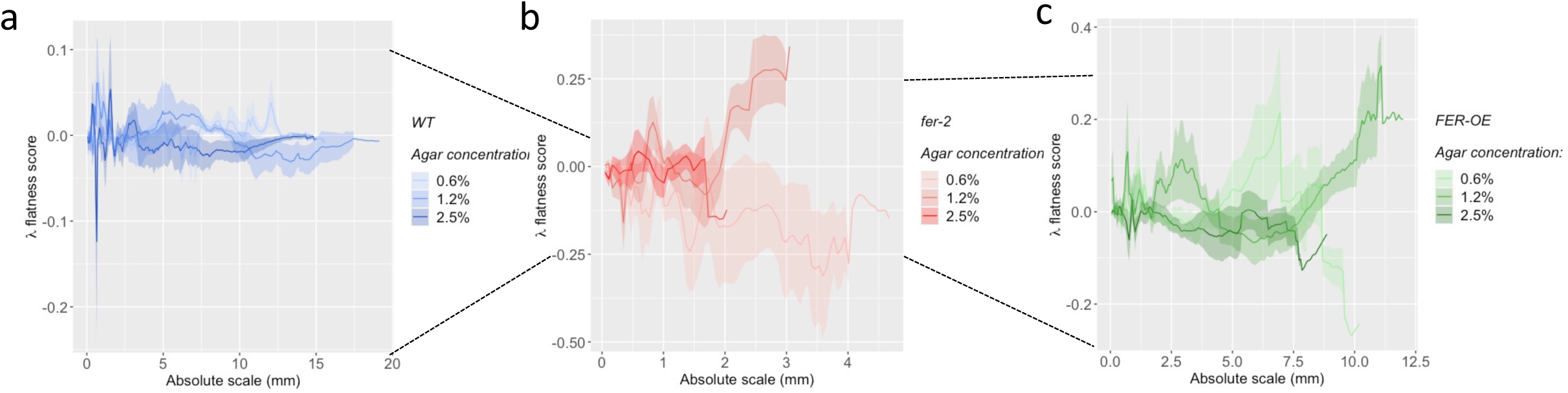
λ flatness score of thalli grown on 0.6, 1.2 and 2.5% agar medium; **a**) λ flatness score of *WT Marhcantia polymorpha*; **b**) λ curvature of the *fer-2* mutants; **c**) λ flatness score of the *FER-OE* transgenics. **a**-**c**) The number of thalli analysed is the same as that of figure 2f, g and h, respectively. The absence of shaded area at high scales indicates that N=1

**Supplementary figure S5:**
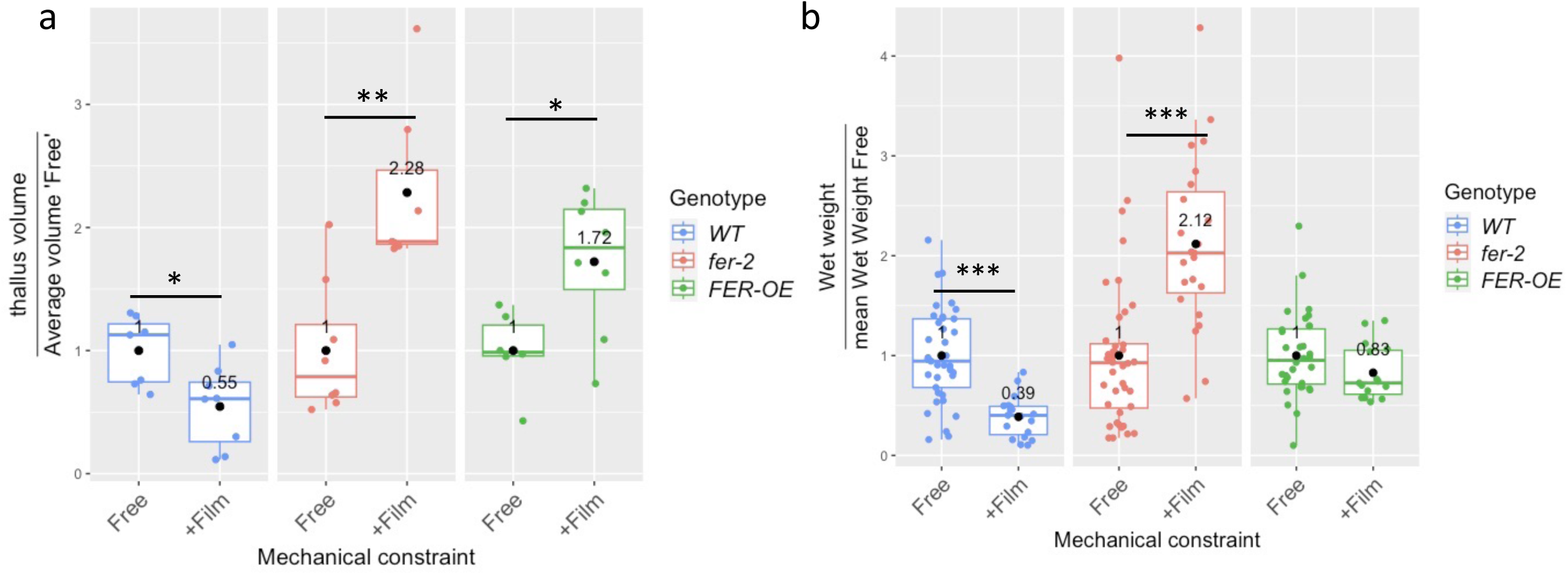
Measurement of volume and thallus weight from thalli grown with and without PDMS films; **a**) Volume of 16-day-old thalli. The number of thalli analysed is the same than that of figure 4e and d; statistics: WT and *fer-2*: Wilcoxon test: * ó 0.05 > p-value ≥ 0.01; **: 0.01 > p-value ≥ 0.001, *FER-OE:* t-test: *: 0.05 > p-value ≥ 0.01; **b**) Weight of 17-day old thalli. t-tests, non-parametric Wilcoxon tests and pairwise tests with a Bonferroni correction were used to assess the relevance of the measured differences. Statistics: WT: a t-test was performed, *fer-2:* a Wilcoxon test was performed. ***: 0.0001 > p-value. Black dots indicate the mean. 37 *WT* thalli without and 19 *WT* thalli with films were analysed, 40 *fer-2* thalli without and 23 *fer-2* thalli with films were analysed and 32 *FER-OE* thalli without and 15 with film were analysed.

**Supplementary figure S6:**
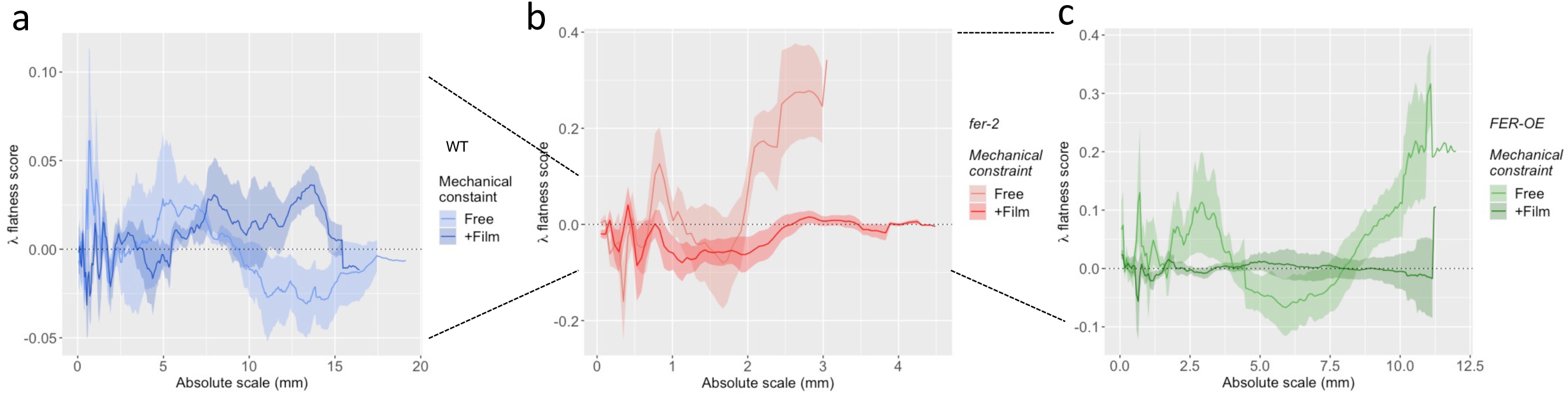
λ flatness score of thalli grown with and without PDMS films; **a**) λ flatness score of *WT Marchantia polymorpha*; **b**) λ flatness score of the *fer-2* mutants; **c**) λ flatness score of the *FER-OE* transgenics. The number of individuals analysed is the same as that of figure 4f, g and h, respectively. The absence of shaded area at high scales indicates that N=1.

